# Complex responses of global insect pests to climate change

**DOI:** 10.1101/425488

**Authors:** Philipp Lehmann, Tea Ammunét, Madeleine Barton, Andrea Battisti, Sanford D. Eigenbrode, Jane Uhd Jepsen, Gregor Kalinkat, Seppo Neuvonen, Pekka Niemelä, Bjørn Økland, John S. Terblanche, Christer Björkman

## Abstract

Insect pests strongly affect the productivity and profitability of agriculture and forestry. Despite the well-known sensitivity of insects to abiotic effects such as temperature, their potential responses to ongoing climate change remain unclear. Here we compile and review documented climate change responses of 31 of the globally most impactful phytophagous insect pests, focussing on species for which long-term, high-quality data are available. Most of the selected species show at least one response affecting their severity as pests, including changes in geographic range, population dynamics, life-history traits, and/or trophic interactions. Of the studied pests, 41% only show responses that are linked to increased pest severity, 4% only show responses of decreased severity, whereas importantly 55%, the majority of studied pests, show mixed responses including both increased and decreased severity under ongoing climate change. Variability in impact is further supported by a thermal sensitivity analysis showing little benefit of climate warming in relation to the optimal developmental temperatures for the majority of these pests under both current climate and future projections. Overall the results show that calculating the net effect of climate change on phytophagous insect pest impact is far from straightforward. The documented variation in responses, e.g. between agricultural and forest pests, indicates that efforts to mitigate undesirable climate change effects must target individual species, taking into account the complex ecological and evolutionary mechanisms underlying their responses.

## Introduction

### Climate change and insect pest impact

Insect pests have major detrimental impacts on agricultural and forestry production^1^ that are likely to increase with anticipated rise in demands for food^2^, bioenergy feedstocks and other agricultural products. For example, animal pests (mainly insects) cause estimated losses of *ca*. 18% of total global annual crop production^3^. Many forest pests, such as the gypsy moth (*Lymantria dispar*) and mountain pine beetle (*Dendroctonus ponderosae*), also have severe ecological impacts: displacing native species, causing widespread defoliation and tree mortality, disrupting ecosystem functions and diminishing biodiversity^4,5^. Further, managing insect pests is generally financially costly. For example, estimated global costs of managing only one pest species, the diamondback moth (*Plutella xylostella*), are 4-5 billion USD annually^6^. Moreover, many agricultural and forest insect pests are also invasive species that contribute to negative ecological consequences and the global costs of managing or mitigating such invasions are estimated to exceed 76.9 billion USD annually^7^.

The substantial global challenges posed by phytophagous insect pests can be exacerbated by ongoing and projected large-scale climatic changes^8^ which could promote increases in pest populations and resulting economic losses^9–12^. Alternatively, pests can be constrained by their environmental niche requirements, physiological tolerances, and phenological or life-history responses to climate, leading to local population declines or extinctions as climates change^13,14^. Clearly, detailed knowledge of insect pests’ current and likely responses to ongoing climate change is essential to counter changing risks. Widespread ecological damage through range expansions and increasing frequencies of outbreaks are increasingly reported^14–17^, but there is a severe deficiency in comprehensive information on insect pests’ responses^18–20^.

### Climate change and insect pest biology

Efforts to predict climate change impacts on insect pests are typically based on empirical studies of distribution responses to geographical and temporal variation in climate, mechanistic studies of physiological responses ^21,22^, mechanistic studies of insect responses to varying abiotic conditions (often in controlled laboratory environments)^23^, climate modelling studies^24,25^, or some combination of these approaches^19^. A common assumption in studies of pests’ responses is that climate-limiting factors are constant across their geographic ranges^26^. Thus studies typically ignore intraspecific variation, a well-known source of variability in climate responses^9,22,27^. Also, pest ranges generally span multiple environments, often including various types of managed landscapes^28^, forming complex dynamic matrices of pest-ecosystem interactions^20,29^. Furthermore, analyses tend to consider a single response (e.g. range expansion), rather than the wide range of pests’ potential responses to climate change^20^, which can be divided into at least four main categories that are non-mutually exclusive^18^: changes in geographic range^30^, life-history traits^31^, population dynamics^32,33^, and trophic interactions^34^ (Fig. 1). Changes in range and particularly population dynamics are likely to be directly linked to economic damage.

To assess current empirically-based knowledge within these four categories of response to climate change, we reviewed primary literature on 31 globally detrimental insect pest species. Species were selected to cover both agricultural and forestry pests, representing various feeding guilds (Fig. S1), being present in various biomes and having large geographic ranges (Fig. 1). Furthermore, we only selected species that have been well studied over a long period. While this approach perhaps leads to biases in terms of geographical range and taxonomy, we feel that it is compensated by having high-quality comprehensive datasets available for the species. This is also critical for allowing an integrated assessment of all the four major response categories outlined above in each species and would not be possible otherwise. As there is a need for more information on biological mechanisms relating to past and present climate change responses in several key biological traits for single organisms^18^, we here provide an update on a number of such mechanisms (range expansion, life-history, population dynamics and trophic interactions) for the selected species in hopes that the data can be used for further predictive modelling. This information is presented in the form of species-specific descriptions and data tables in Supplement 1. We also identify critical knowledge gaps, and highlight aspects that require further research to anticipate, mitigate and manage climate-driven changes in pest impacts.

**Fig. 1.**
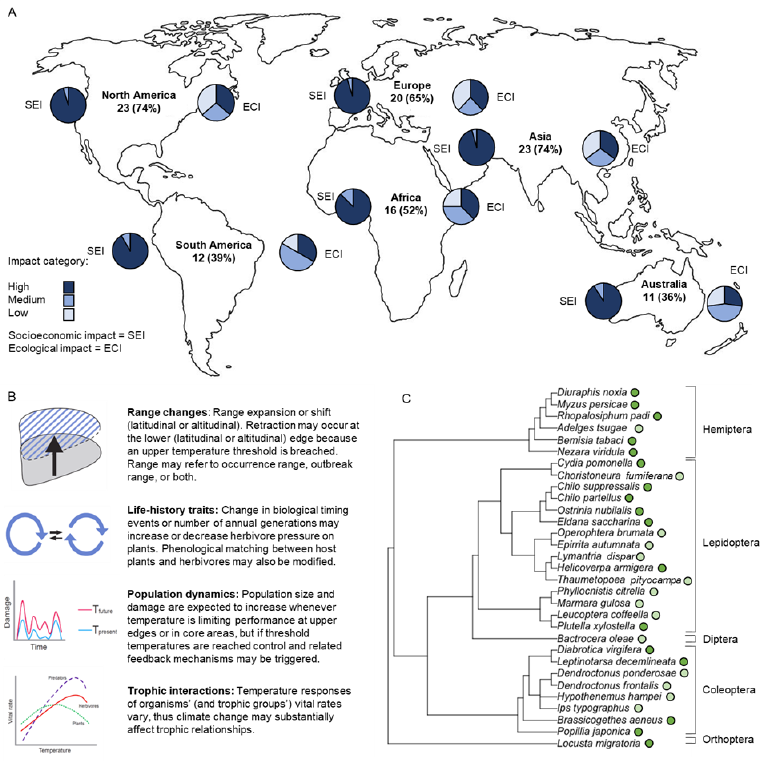
The distribution of 31 insect pests according to (A) the number of species in the study occurring in each continent (with % of all those included) according to CABI. Note that many species occur on multiple continents. Flanking each continent are pie charts showing the distribution of socioeconomic impacts and ecological impacts caused by these species. (B) Schematic representation of four major categories of responses to climate change: range changes, life-history traits, population dynamics and trophic interactions (see Supplement 2). (C) A phylogenetic tree (compiled from the Tree-of-life project) of the 31 species considered in this analysis. Dark green circles reflect pests on annual crops (mainly agricultural pests) and light green circles pests on perennial crops (mainly forestry pests).

## Materials and methods

### Data selection

Thirty-one of the socioeconomically and ecologically most detrimental phytophagous insect pests globally were selected that collectively: infest both agricultural and forestry crops, represent diverse feeding guilds, originate from both tropical and temperate environments, have large geographic ranges (preferably covering several continents), and have been well studied and monitored over recent decades (Fig. 1). A lack of rigorous long-term monitoring, with consistent sampling effort, is probably the biggest limitation hindering efforts to characterize biological systems’ responses to climate change robustly. Because of their large economic impact, insect pests represent a group of organisms with relatively good data compared to other groups; data are collected frequently but not consistently and data quality tends to be positively correlated to density and range expansion of the species. Thus, pests are good models for such efforts because abundant information about their distributions, impacts and interactions is routinely collected. However, since we selected species with large ranges, our results can be biased towards responses of species with broad thermal niches, thus the indicated general effects of climate change are likely conservative. Further, since habitats strongly affect insect ecology, we assume that species in disparate habitats will have different potential responses to climate change, so we chose species prevalent in a wide spectrum of lightly-managed to heavily-managed habitats. Then, using Web of Science searches (Thomson Reuters), we selected three types of studies. First, studies that compared climate trends and empirically determined trends in relevant aspects of the chosen pests, e.g. range, abundance or damage (economic and/or ecological). Second, studies that tracked population-dependent differences in relevant traits (e.g. voltinism) of the pests across time. Third, studies that modelled attributes of the pests, including a substantial historical data component. Data sources include studies published in scientific journals, pest management databases (e.g. EPPO and CABI) and records from national environment/pest management institutions. We also contacted several experts for assessments of data quality. The short summaries describing each pest species can be found in Supplementary File 1.The responses recorded in these studies were classified into four major types (Fig. 1B), and as either increasing or decreasing pest severity (Table S1). We used a modified version of a generic impact score system to assess impact and severity^35^. The impact criteria can be found in Supplement 2 and the qualitatively assigned categories are found in the attached datafile. As has been suggested in several recent studies^10,18,36^ holistic integrated analyses are to be preferred over single-trait analyses when assessing climate change responses, and this is what we attempted to achieve with our approach. Thus while the present study is neither a formal meta-analysis nor exhaustive, it synthesizes current knowledge of integrated climatic responses of 31 pests with the aim to illustrate general patterns, problems and challenges, in a precautionary manner.

### Rank order correlation

Associations between explanatory and response variables regarding effects of climate change on the 31 selected serious insect pests were explored by Kendall rank order correlation analysis. The results are presented in Table S2, and the following list explains abbreviations and the range of these variables, which are listed in the beginning of Supplement 2. NRT = Number of response categories (1 – 4), PA = Perennial or annual host (1 [perennial] – 3 [annual]), IE = Internal or external feeder (1 [external] – 2 [internal]), BRANK = Mean habitat biome ranked from tundra to tropical (1 [tundra] – 4 [tropical]), AF = Agricultural or Forestry pest (1 [agricultural] – 2 [forestry]), SEI = Socioeconomic impact (1 [low] – 3 [high]), SEId = Change in Socioeconomic impact (1 [decrease] – 3 [increase]), ECI = Ecological impact (1 [low] – 3 [high]), ECId = Change in ecological impact (1 [decrease] – 3 [increase]), GD = Difference in responses to climate change between geographical areas of range (1 [no] – 2 [yes]). This analysis was run in SPSS v. 24.0 (IBM Corp., Armonk, NY, USA).

### Optimal temperature in the past, the present and the future

A meta-analysis on optimal temperatures of the 31 insect pest species was conducted to quantify potential climate change stress. We extracted optimal temperatures for development (T_opt_) for the species from the primary literature, giving priority to studies investigating temperature dependence of the whole life-cycle, as well as using populations from the core of the range (Table S3). Latitude and longitude coordinates were either copied straight from the article, or extracted from global maps based on the sampling location reported in the original article.

Ambient temperatures at each location in our species database (Table S3) were extracted from a Global Circulation Model that forms part of the *Coupled Model Intercomparison Project* phase 5^37,38^, which we sourced directly from the Earth System Grid database (http://pcmdi9.llnl.gov/). More specifically we considered predictions of average monthly near surface temperature (ambient temperature hereafter, T_amb_) from the HadGEM2-CC model^39^. For the present and future conditions, we considered models with a radiative forcing of 8.5Wm^−2^ (Representative Concentration Pathway 8.5), the most extreme climate warming scenario included in the IPCC Fourth Assessment report^8^, and that which is most representative of current trajectories^40^. Here, we aimed to capture “present” ambient temperatures (2006-2015), “near-future” ambient temperatures (2056-2065) and “future” ambient temperatures (2070-2079). The “past” ambient temperatures (1960-1969) were extracted from the historical experiment of the same model. Across each of these four decades, we calculated an overall average mean temperature from the 12 monthly averages for each year. As species at high latitudes in the northern hemisphere undergo a period of dormancy during winter (and hence are buffered from winter temperatures), for locations above 45° latitude (15 of 38 locations, Table S3), we considered only temperatures during the summer months from May to September inclusive. Data were extracted from raw files, and subsequently cleaned using functions in the “raster” package for R^41^. The full R-code workflow can be found at GitHub: [https://github.com/madeleine-barton/Complex_pest_responses].

The overall T_amb_ for each of the time periods were compared against the species T_opt_ at each location in two ways. First by visually comparing the differential between T_opt_ and T_amb_ (Fig. 3), where a small value (close to 0) indicates high thermal suitability, and then with a phylogenetically corrected generalized linear least square model (pgls) investigating the relationship between thermal suitability (expressed as T_amb_/T_opt_) and absolute latitude. A high value (close to 1) indicates high thermal suitability. Models were run using primarily the “pgls” function in the “caper” package for R^42^. Overall model results are shown in Table S4 and the full R-code workflow can be found at GitHub: [https://github.com/madeleine-barton/Complex_pest_responses].

## Results and discussion

### Insect pest responses to contemporary climate change are complex

Of the 31 insect pest species selected for the study, 29 (94%) reportedly show some response attributable to contemporary climate change (Table S1), and 28 (90%) present more than one response (Fig. 2a). Of the 29 showing some response 26 (90%), 18 (62%), 16 (55%) and 4 (14%) respectively show changes in: geographic range, population dynamics, life-history (traits related to phenology and voltinism), and trophic interactions (Fig. 2b). While at least one reported response of almost all of these species is likely to increase pest severity (e.g. range expansion or increases in population density), 59% (17/29) of them also show responses likely to reduce pest severity (e.g. range contraction or decreased physiological performance), and often this reduction occurs simultaneously with other responses likely to increase severity (Fig. 2c). The most common severity-reducing responses are reduction in pest population density (13/29), followed by range contraction (6/29) (Fig. 2c).

**Fig. 2.**
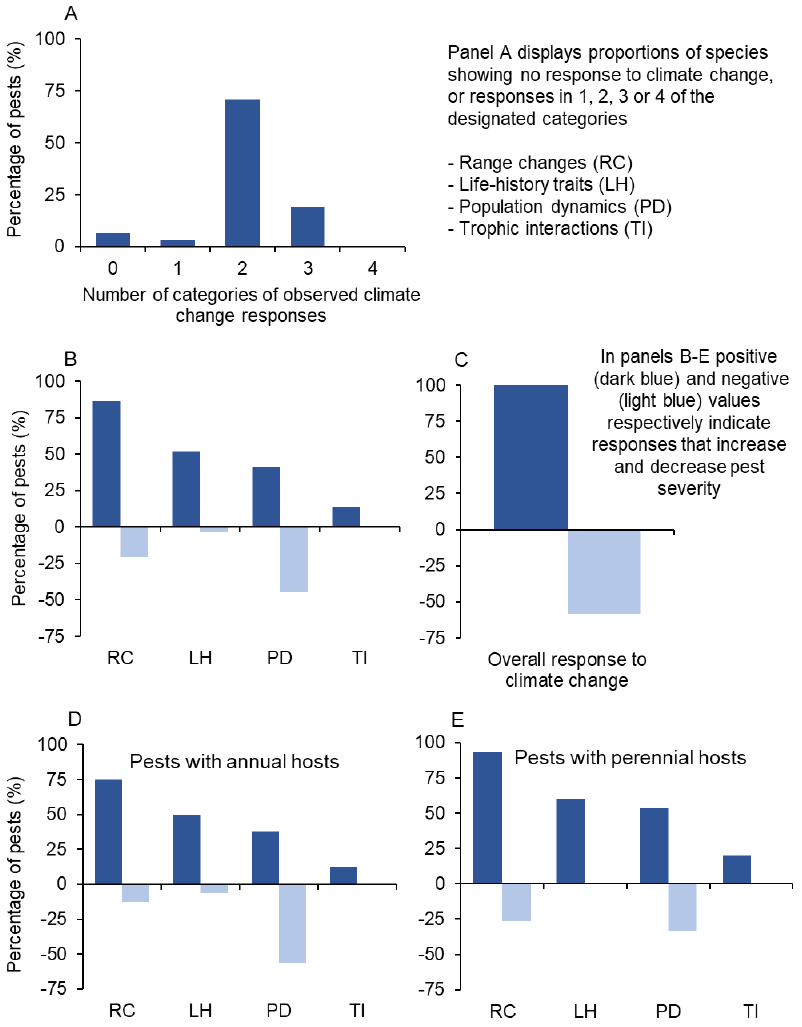
Responses to climate change of 31 insect pests with high socioeconomic and/or ecological impact. (A) Shows the number of species responding in 0 to 4 traits to ongoing climate change. Dark and light blue columns in (B-E) show percentages of the 31 insect pest species displaying severity-increasing responses (e.g. increased range) and severity-decreasing responses (e.g. decreased economic damage due to smaller population size) to climate change in the four traits investigated here. Single species may show responses to multiple and (B-E) only display data for the 29 species that showed some response attributable to climate change (see Supplement 2). Observe that in (B-E) some species showed no response in some traits, so total percentages in these cases are less than 100% (i.e. if all 29 species show a response increasing severity due to range expansion, this trait would receive a value of 100%).

Responses of 59% (17/29) of the pest species with reported sensitivity to contemporary climate change have also varied between different parts of their ranges. For example, the range of the Colorado potato beetle (*Leptinotarsa decemlineata*) has expanded northwards in recent decades, and its population density has increased in core European areas (Table S1). The range of the winter moth (*Operophtera brumata*) has also expanded, towards higher latitudes and more continental areas at the northern European edge of its range, and its trophic interactions have changed in the boreal-tundra ecotone, where outbreaks have spread from the main host *Betula pubescens* to an alternative host (*B. nana*) above the tree-line (Table S1). Several species also show both severity-increasing and severity-reducing responses in different parts of their ranges. Notably, thermal tracking^43,44^ has been observed in some species (4/17), e.g. the spruce budworm (*Choristoneura fumiferana*; Table S1) has expanded its geographic range towards higher latitudes while it has retracted, or its abundance has declined, at lower latitudes. Similarly, northward range expansion of the hemlock woolly adelgid (*Adelges tsugae*) has been observed in the USA, while the economic damage it causes is decreasing in the southern part of its range due to poor heat tolerance of young nymphs during summer (Table S1).

### Do responses of phytophagous pests on annual and perennial crops differ?

The main response patterns of pests of annual (mainly agricultural pests) and perennial (mainly forestry pests) crops are similar, with some subtle differences. Contrary to expectations based on differences in feeding or host ecology, and evolutionary constraints, pests of annual crops show more severity-reducing responses than pests of perennial crops (e.g. trees). To assess the potential impact of agricultural and forest pest responses to climate change, we categorized the species according to their historic and current socio-economic and ecological impacts, and effects of contemporary climate change on those impacts. Overall socio-economic and ecological impacts have reportedly increased across the geographic ranges of species that have responded to climate change^11,20,20^. More importantly, while all the considered pests on perennial crops already have large ecological impact, 85% (17/20) of the pests on annual crops currently have relatively low ecological impact beyond the cropping systems they infest. However, climate change might be inducing increases in the relatively low impact of some pests on annual crops. For instance, the green stink bug (*Nezara viridula*) and maize stem borer (*Chilo partellus*) displace native bugs and borers, respectively, as their ranges expand (Table S1). Further, the range of the western corn rootworm (*Diabrotica virgifera virgifera*) in Europe has expanded, and it can cause large ecological damage by spreading maize chlorotic mottle virus to several natural hosts (Table S1). A potential explanation is that reductions in phenological constraints associated with climate warming (mediated for instance by increases in host growth season, or shorter and milder winters^45^), can increase interactions between pests in annual agricultural habitats and surrounding ecosystems^36,46^, thereby increasing ecological impacts. Indeed even small phenological mismatches might have large knock on effects for ecosystem function and predator prey interactions^14,36^.

In addition to the fact that latitudinal differences in pest distributions might modulate climate change effects, several other mechanisms could be involved in the divergence of responses in annual and perennial systems. Unlike forestry pests, agricultural pests are generally associated with fragmented habitats^47^ and may therefore have higher local extinction risks due to Allee effects when climate changes^13^. Further, while climate change can disrupt biological control by natural enemies in either annual or perennial systems^48^, the biological control agents frequently introduced in annual systems may have lower genetic diversity than native agents, and hence lower adaptive capacity to respond to environmental changes^49^. Direct effects of climate change on the performance and phenology of pests have been detected in both annual and perennial systems. Since pests often persist through part of the season in a resting or dormant stage, especially at high latitudes and/or altitudes^45^, climate change can contribute to phenological mismatches between hosts and emergence of key life-stages^14,22,46^, as seen in *O. brumata* (Table S1). However, pests in annual and perennial systems might differ in general susceptibility to phenological mismatching, *inter alia* the former might be more sensitive to phenological host limitation; especially relative to bark beetles and root feeders. Taken together, while there are some differences that seem to associate with whether the system is annual or perennial, pests in both systems show large variability in how ongoing climate change is affecting both their ecological and socioeconomic impact.

### Past, present and future temperature stress on the major insect pests

It has been argued that pests may suffer negative consequences of ongoing climate change owing to reduced thermal suitability and increasing frequency of high temperature extremes leading to population reductions^50^. For further exploration of this in our focal species, we assess the proximity of optimum development temperature (T_opt_) of the 31 pest insects compared to their ambient habitat air temperatures (T_amb_) (Fig. 3). Relating ambient temperature during the growing season in past, present and future climates to T_opt_ shows large variability in how pests are expected to benefit from climate change owing to regional complexity. In general, warming climates are expected to be beneficial for growth and development, and indeed, in all but two cases T_amb_ closely approached T_opt_ when comparing past, current, near future and future climates (Fig. 3B). This conclusion was also supported by a phylogenetically-informed regression analysis (Table S4). Further, this analysis suggested that pests at higher latitudes have greater disparity between T_amb_ and T_opt_, indicating greater capacity to benefit from climate warming, unlike more low latitude pests that are already close to T_opt_. Low latitude species also potentially risk increasing frequency and intensity of heat stress as climate warms^51^, a notion receiving support in a recent analysis of the upper thermal tolerance of 15 dipteran pests^50^.

**Fig. 3.**
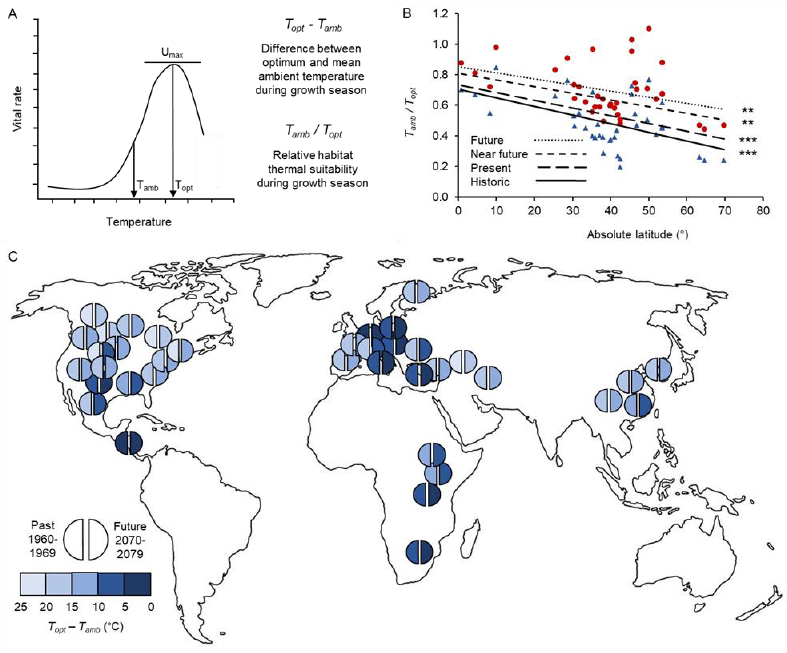
Summary figure of thermal sensitivity analysis of 31 insect pests. As input we use published optimum temperatures of the species (T_opt_, the temperature at which performance is maximised, Umax) and mean ambient temperature (T_amb_) during the growing season. This includes the whole year below 45°S/N, and the summer months above 45°S/N. (A) Schematic thermal performance curve including the two metrics extracted. (B) Here T_amb_ / T_opt_ is plotted against latitude for the four periods investigated (historical: 1960-1969 [blue triangles and dotted line], present: 2006-2015 [fine dashed line], near future: 2056-2065 [coarse dashed line] and future: 2070-2079 [red circles and *solid line]). Stars denote significant correlations in a phylogenetically corrected generalized linear least square model: * = P<0.05, ** = P<0.005. (C) Shows how many degrees T_amb_ differs from T_opt_ in past (left half of circle) and future (right half of the circle) climates. Circles have been placed in the approximate location where individual studies sampled the respective pests. Darker colors reflect ambient temperatures near the optimum temperature and therefore climates likely beneficial for pests.*

However, examination of patterns in more species, as well as on other thermal traits, especially upper thermal limits or feeding (damage) rates, would be required to validate this hypothesis. Agricultural pests accounted for only 4% of the ca. 380 species included in the database of upper thermal limits compiled by Hoffmann et al.^52^, highlighting a potential information gap in the current literature. While the pests in the current data represent a wide geographic distribution (Fig. 1A), the studies on T_opt_ used here mostly reflect populations sampled in the northern hemisphere (Fig. 3C). This is a general problem found in other large-scale analyses of climate change responses, such as phenology^36^ and insect metabolic or development rate-temperature databases^53^ showing a need for further studies covering underrepresented locations. Finally, as air temperatures are reported in the global temperature database, there is risk of underestimation of microclimate variability^51^ and thus the extent of potential buffering owing to three-dimensional habitat complexity of operative temperatures^51,51,54^.

### Evolutionary responses of insect pests to climate change

Insect pests may evolve rapidly in response to contemporary climate change^16,55–59^. Thus, apparently sound projections of insect pest responses to climate change^11^ may be compromised if evolutionary responses are not considered^60^. Indeed, rapid evolutionary effects have influenced - or could influence further - projections for several of the 31 species considered here (see Supplement 1). For example, disruption of phenological synchrony between *O. brumata* and oak in temperate Europe due to increasing temperatures^30^ has been apparently restored by a hereditary change in egg hatching dates^61^. Also, range expansions of some of the forestry pests induced by climate change have resulted in colonization of areas with novel host tree species that have little innate resistance due to lack of co-evolution with the pests^5^. In contrast, the similarity of crops grown across large areas might promote co-evolution between agricultural pests and their hosts^62^. Links between biological invasions or range expansion events, climate change and evolutionary processes have received recent attention^9,17,21,59^, but there is still pressing need for further research in this field. The effects of management practices and evolution have generally been considered too much in isolation, especially in climate-change contexts^18,49^.

## Conclusions

The 31 widely-distributed pest insects that seriously affect agricultural or forestry systems studied here show multiple and varying responses to climate change. By providing an up-to-date database that reviews biological responses to climate change in the selected pests (Supplement 1) we offer standardized information that can be further explored by other researchers. Although the present analyses cannot be considered absolute, complete, and without taxonomic, geographic and study intensity biases^10^, we nevertheless detected several overarching patterns that allow us to draw some general conclusions.

1. The data suggest that determining the net severity change of pests to climate change is complex since most species considered here have shown multiple responses that vary spatially^24^. The present study also provides evidence for mixed directionality of responses as well as potential explanations thereof based on general mechanisms. This set of complex but predictable outcomes and regional heterogeneity of responses is challenging for management but cannot be ignored as it is the emerging consensus in this and other studies^11,19^.
2. The current study urges caution in performing large-scale analyses only with single traits, since single pests often show mixed directionality of effects of climate change in different traits. Lacking the interactions among different traits in each pest species may easily lead to incomplete conclusions. To correct this we recommend more indepth studies of biological mechanisms in a few representative species. For example, a recent meta-analysis shows that models integrating biological mechanisms from multiple traits significantly improve predictions of climate change impacts on global biodiversity^18^.
3. Mounting evidence suggests that pests and their hosts are responding not only through ecological, but also evolutionary processes to climate change^17,57,59^. Thus, evolutionary approaches might be under-exploited in pest management strategies^49^. Including evolutionary and ecological information when formulating integrated management strategies may facilitate robust intervention and control (as recently demonstrated in disease vector control programs^63^). Furthermore, it would be useful to pinpoint species with high evolvability in traits relevant to climate change^17^, or that show trade-offs between traits linked to basal climatic stress resistance and plasticity^59,64^.
4. Combining data from large-scale experiments (e.g. mesocosm) and computational models may improve estimates of climate change effects^19,59,65^. Experiments should be designed to assess variance components with indicated importance in climate modelling studies, to identify the factors related to climate change that most strongly influence pest population growth and performance, such as for example the increased feeding efficacy of the Japanese beetle (*Popillia japonica*) on carbon dioxide-enriched soybean^66^. Indications that the response to climate change differ among trophic levels, translating into shifts in the relative importance of bottom-up and top-down population processes^67^ needs to be studied further as even relatively small changes could result in large effects when multiple interactions are affected simultaneously^68^. Standardized experiments enable high-throughput investigation of pests (for recent example see^69^) and facilitate the development of watchlists or prioritization tools (such as The UK Plant Health Risk Register^70^) of key species that require further study. However, as the current data suggest large regional variability in pest responses to climate change, national or regional databases, while excellent locally, might offer poor insight into invasions into other regions unless coordinated or standardized efforts are attained, especially across political boundaries.
5. As T_amb_ is generally increasing towards T_opt_ for growth and development in these species, there is an expectation of increasing pest severity under future climate scenarios^71^. However, the relative benefit of increasing ambient temperatures is negligible for many of the studied pests (Fig. 3C). Indeed, since low-latitude species already showed T_amb_ close to T_opt_, as climates warm T_amb_ for these species may surpass T_opt_, thus decreasing pest severity, under future climates^50,51^.
6. Finally, and importantly, the patterns of regional variability and complexity described here are likely to apply to non-pest insects as well as non-insect species in addition to the 31 insect pest species assessed here. The extent of generality of responses across various taxa will be important to assess in future studies^14,20,59,65^.

## Acknowledgements

The authors thank Christer Wiklund, Stig Larsson and Myron Zalucki for insightful comments, and all contributors to the book, “Climate Change and Insect Pests” edited by C. Björkman and P. Niemelä, published in 2015 by CABI publishing. The work was financially supported by the research program ‘Future Forests’. GK acknowledges financial support from the Leibniz Competition (SAW-2013-IGB-2). SDE acknowledges financial support by the US Department of Agriculture’s National Institute of Food and Agriculture (award #2011-68002-3019).

## Author contributions

All authors jointly designed the study and collected species data. SN performed the rank correlation analysis, PL, JST and MB performed the optimum temperature analysis. All authors contributed to preparation of the supplements. PL, MB, AB, SDE, JST and CB prepared the first draft of the paper, and all authors edited the final version. The authors declare no conflicts of interest.

**Supplement 1:** Species summaries

**Supplement 2:** Extended materials and methods

